# Conductive particles enable syntrophic acetate oxidation between Geobacter and Methanosarcina from coastal sediments

**DOI:** 10.1101/213843

**Authors:** Amelia-Elena Rotaru, Hryhoriy Stryhanyuk, Federica Calabrese, Florin Musat, Pravin Malla Shrestha, Hannah Sophia Weber, Oona L.O. Snoeyenbos-West, Per O.J. Hall, Hans H. Richnow, Niculina Musat, Bo Thamdrup

**Author notes:** **Correspondence**: 1. **Amelia-Elena Rotaru**, PhD, Assist. Prof. Department of Biology University of Southern Denmark Address: Campusvej 55, 5230, Odense M, DK. **Correspondence**: 2. **Niculina Musat**, Dr., Scientific Head ProVIS - Centre for Chemical Microscopy Department of Isotope Biogeochemistry Helmholtz Centre for Environmental Research Address: Permoserstraße 15, 04318 Leipzig, Germany.

## Abstract

Coastal sediments are rich in conductive minerals, which could impact microbial processes for which acetate is a central intermediate. In the methanogenic zone, acetate is consumed by methanogens and/or syntrophic acetate oxidizing (SAO) consortia. SAO consortia live under extreme thermodynamic pressure and their survival depends on successful partnership. Here we demonstrate that conductive minerals facilitate a SAO partnership between *Geobacter* and *Methanosarcina* from the coastal sediments of the Bothnian Bay, Baltic Sea. Bothnian methanogenic sediments showed a high apparent isotopic fractionation (α_c_ 1.07) characteristic of CO_2_-reductive methanogenesis. The native community was represented by electrogens such as *Geobacter* and methanogens like *Methanosarcina*. Upon the addition of conductive particles (activated carbon and magnetite) methanogenesis from acetate increased four fold. *Geobacter* (96% related to *G. psychrophilus*) and *Methanosarcina* (99% related to *M. subterranea*) dominated the conductive particle-spiked SAO communities. Using NanoSIMS we demonstrated that during SAO, *Geobacter* incorporated 82% of the labeled acetate as compared to only 18% by *Methanosarcina*. At the same time *Geobacter* converted 27% of the ^13^C-acetate to ^13^CO_2_ as detected by IRMS. Indigenous soluble shuttles were not involved in SAO, since spiking fresh cultures with spent-media filtrate had no effect on methanogenic rates. Our results demonstrate that *Geobacter* oxidizes acetate to CO_2_ while transferring electrons extracellularly via conductive particles to *Methanosarcina*, which utilizes them for CO_2_ reduction to methane. In natural environments, mediation of SAO by conductive particles between electrogens and methanogens could impact the iron and methane cycles.

**Significance:** Acetate oxidizing bacteria are known to thrive in mutualistic consortia in which H_2_ or formate is shuttled to a methane-producing *Archaea* partner. Here we discovered that they could instead transfer electrons via conductive minerals. Mineral-SAO (syntrophic acetate oxidation) could be a vital pathway for CO_2_-reductive methanogenesis in the environment, especially in sediments rich in conductive minerals. Mineral-SAO is therefore of potential importance for both iron and methane cycles in sediments and soils. Additionally, our observations imply that agricultural runoff or amendments with conductive chars could trigger a significant increase in methane emissions.

## Introduction

Syntrophic acetate oxidizing (SAO) bacteria live in a mutualistic interaction with methanogenic archaea which feed on the H_2_ or formate released by the SAO bacterial partner (1). Besides H_2_ or formate, cysteine can also be used to transfer electrons in some SAO consortia (2). Several studies with synthetic consortia have shown that SAO can be performed by members of the phylum *Firmicutes* (*Thermacetogenium, Clostridium, Thermotoga*, candidatus*‘*Contubernalis’ and *Syntrophaceticus*) and *Proteobacteria* (*Desulfomicrobium* and *Geobacter)* (2–14). Remarkably, acetoclastic methanogens (*Methanosarcina* and *Methanothrix*) were proposed to play the role of syntrophic acetate oxidizers when provided with an appropriate H_2_-consuming partner (15, 16). Some of the genera above were suggested to carry out SAO in thermophilic digesters (17–26), lake/river sediments (21, 27, 28), tropical wetland soil (29), rice paddies (30–32), or oil field reservoirs (33). Many of these environments are rich in (semi)conductive minerals like magnetite (34, 35), pyrite (36, 37) or black carbon resulting from incomplete burning of plant biomass (38–40). Electrically conductive iron-oxide minerals and carbon chars (magnetite, granular activated carbon, biochar) were previously shown to stimulate **d**irect **i**nterspecies **e**lectron **t**ransfer (DIET), a recently described form of interspecies electron transfer (12, 41–49), whereas strict H_2_-based interactions were shown to remain unaffected by the addition of conductive materials (44). DIET is a syntrophic association where electrons are transferred via conductive and/or redox active cell surface structures between an electron-donating species (electrogen) and an electron-accepting species (electrotroph) (47–49). Conductive minerals seem to alleviate the need for cells to produce certain cell surface molecules required for DIET (41). DIET mediated by conductive materials is considered a novel strategy to stimulate recalcitrant organic matter decomposition in anaerobic digesters (50–52) and to enhance methanogenic decomposition of organics in rice paddies (46, 53) and aquatic sediments (28, 54). It is likely that conductive materials replace the molecular conduits that cells require to establish direct contacts during DIET.

Although SAO via DIET was considered thermodynamically favorable at pH values between 1.9 and 2.9, and impossible at pH 7 (55), conductive minerals were shown to facilitate SAO in synthetic denitrifying consortia at pH 7 (56). Nevertheless, the impact of minerals on environmentally relevant SAO is presently not understood. Mineral facilitated SAO could be significant in coastal environments rich in (semi)conductive minerals (36, 57–59). Such (semi)conductive minerals are likely to impact microbial processes (36, 56) for which acetate is a central intermediate (60–63).

Here we investigated the role of mineral-SAO in methanogenic processes from coastal sediments. We examined if electrically conductive materials mediate SAO between *Geobacter* and *Methanosarcina* co-existing in the brackish, iron rich coastal sediments of the Bothnian Bay. Our results indicate that mineral-SAO may impact both the iron and the methane cycle in these sediments, with implication for atmospheric methane emissions.

## Results and discussions

In this study we show that methanogenic communities from the Bothnian Bay made use of (semi)conductive particles to facilitate SAO. For this we used a combination of physiological and stable isotope labeling-experiments followed by monitoring of labeled products and incorporation of the labeled-substrate in phylogenetically assigned cells by Nanoscale Secondary Ion Mass Spectrometry (NanoSIMS) coupled with CAtalyzed Reporter Deposition Fluorescent In Situ Hybridization (CARD FISH).

Syntrophic acetate oxidizers are difficult to enrich (57) because SAO is thermodynamically challenging (55). Here, we have successfully enriched SAO consortia from temperate sediments (sediment temperature 15°C, incubation temperature 20–25°C) by successive cultivation in the presence of electrically conductive (>1000 S/m (58)) granular activated carbon (GAC).

### Characteristics of the Bothnian Bay methanogenic zone

#### Biogeochemistry

Our hypothesis was that a high conductive mineral content would stimulate electric interactions between abundant electroactive microorganisms co-existing in the methanogenic zone. The Bothnian Bay sediments are rich in conductive minerals either dispersed within the fine structure of sediments or within ferromanganese nodules (59).

To explore mineral-mediated interactions in the Bothnian Bay, we sampled the methanogenic zone of these sediments to verify the mineral content. Sediment cores were collected from 15m water depth at station RA2 located at 65°43.6′N and 22°26.8′E in the Bothnian Bay (Fig. 1), which had high sediment temperatures (15°C) and low *in situ* salinity (0.5). The mineral content was low in insoluble manganese oxides (13±3 µmol/cm^3^ from both HCl and dithionite extractions), high in insoluble FeS (HCl-extractable 229±8 µmol/cm^3^), high in crystalline iron oxides such as semiconductive goethite (dithionite extractable Fe^2+^ and Fe^3+^ 131±4 µmol/cm^3^) or conductive magnetite (32±7 µmol/cm^3^ oxalate-extractable). Magnetite content was similar to what has been previously observed below the sulfate-methane transition zone in Baltic Sea sediments (circa 30 µmol/cm^3^) (60).

**Figure 1.**
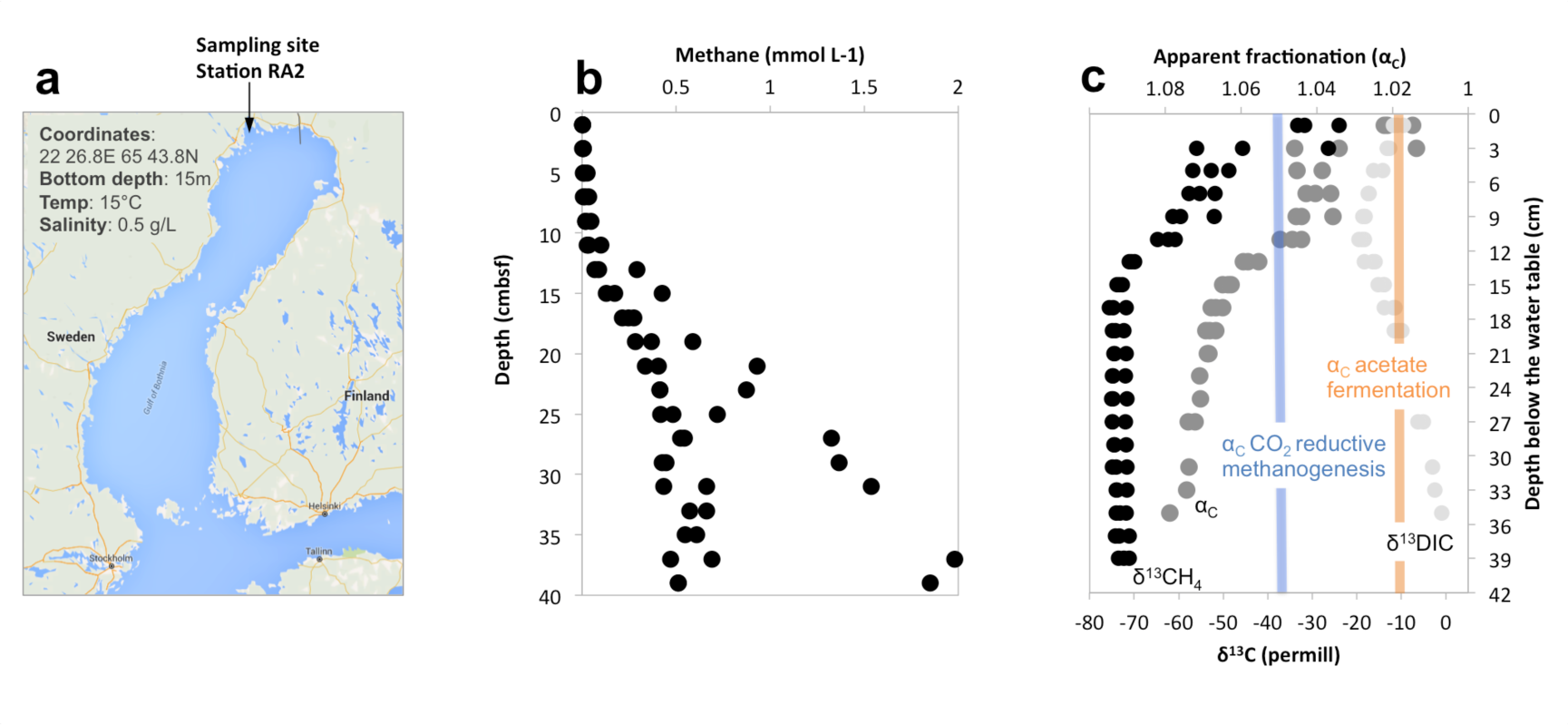
CO_2_ reductive methanogenesis in the Bothnian Bay methanogenic zone. The sampling site, RA2 was off the Bothnian Bay northern coast (a). Here, methane accumulated close to and sometimes over the saturation limit (b) and was strongly depleted in ^13^C (low δ^13^CH_4_), which indicated a high apparent fractionation (α_C_) characteristic of CO_2_-reductive methanogenesis (c). Previous studies showed a α_C_ of ca. 1.05 (blue line) in *Methanosarcina* grown via CO_2_-reductive methanogenesis (83–85). A α_C_ of ca. 1.02 (orange line) was observed in *Methanosarcina* species grown by acetoclastic methanogenesis (86, 84).

Besides iron-oxide minerals, previous studies showed that black carbon, also a conductive material (40), dominated the coastal sediments of the Baltic Sea representing 1.7% to 46% of the total organic carbon (TOC) in sediments closer to coastal towns (61). Conductive materials could reach the Bothnian Bay by river runoff from the eight rivers entering the bay from Sweden and Finland, but also runoff from forestry and various coastal industries (59, 62).

The high abundance of conductive particles is likely to stimulate electrical interactions between abundant electroactive microorganisms co-existing in the methanogenic zone (41–43, 45, 52). Methane reached its highest concentrations below 25 cm depth (Fig. 1). In the methanogenic zone two independent processes, SAO and/or acetoclastic methanogenesis could consume acetate, a key intermediate of organic matter decomposition. SAO bacteria would need a CO_2_-reductive methanogenic partner to scavenge the electrons released during acetate oxidation. To find out if CO_2_-reductive methanogenesis was occurring in these sediments, we looked at the apparent isotopic fractionation of dissolved organic carbon (DIC) and methane. Methane was strongly depleted in ^13^C relative to DIC (median δ^13^CH_4_ of - 74‰ and median δ^13^DIC of −2.5‰) (Fig. 1), which results in a signature apparent isotopic fractionation (α_c_) of 1.07, characteristic of CO_2_-reductive methanogenesis (63).

#### Microbial community

DIET consortia (*Geobacter* and *Methanosarcina*) can usually form more efficient electron transfer associations via conductive minerals than they do in their absence (42–44, 64). In contrast, H_2_-transfering consortia remain unaffected by conductive materials (44). We predicted that Bothnian Bay sediments rich in conductive minerals are favorable for mineral DIET-associations. As anticipated, these iron mineral-rich sediments harbored *Proteobacteria* including exoelectrogens related to *Geobacter* and *Rhodoferax*, and *Archaea* methanogens related to *Methanosarcina* (Fig. 2 and Suppl.file.1). Both *Geobacter* and *Rhodoferax* were previously shown to form DIET-associations with species of *Methanosarcinales* (48, 64, Rotaru & Lovley, unpublished). Until now, only *Methanosarcinales* were shown to establish DIET associations with electrogens (48, 49, 64), probably due to their high c-type cytochrome content, which allows for electron uptake from electrogens (48, 65).

**Figure 2.**
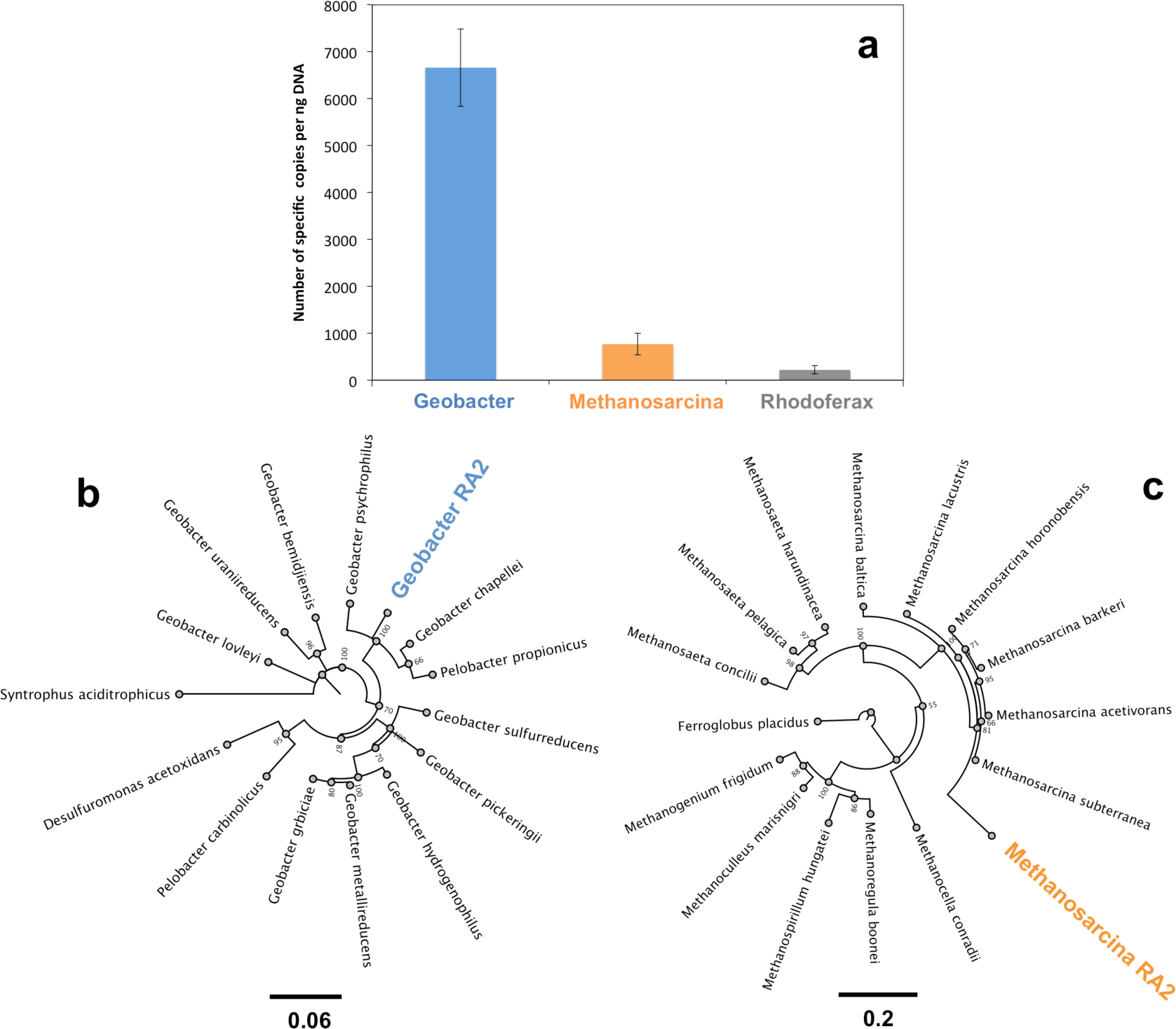
Microbial community in the Bothnian Bay methanogenic zone, featuring microorganism associated with DIET. Quantitative PCR (a) showed that *Geobacter* were the most abundant electrogens, and we could also detect *Rhodoferrax*. No other electrogens could be detected. DIET-associated *Methanosarcina* were the only detected methanogens (a). Maximum likelihood tree of the 16S rRNA gene of the exoelectrogen, 96% related to *Geobacter psychrophilus*, enriched during SAO mediated by activated carbon (b). Maximum likelihood tree of the 16S rRNA gene of the Methanosarcina, 96% related to *M. subterranes*, enriched during SAO mediated by activated carbon (c).

Based on the observations that i) sediments are high in conductive mineral content; ii) CO_2_ reductive methanogenesis prevailed, and iii) *Methanosarcina* and electrogens cohabited, we anticipated that mineral DIET could occur in the methanogenic zone of the Bothnian Bay. We tested this hypothesis in sediment incubations with or without the addition of exogenous conductive particles.

GAC facilitated methane production from acetate (Fig. 3a) and other substrates (ethanol, butyrate and glucose) that were degraded via acetate (Suppl.file.1). GAC was the preferred conductive particle because we could concentrate rigorously on electron transfer (42), whereas by using (semi)conductive magnetite (Fe^II^Fe^III^_2_O_4_) its Fe^III^ content would additionally drive iron reduction especially during long-term incubations (66).

**Figure 3.**
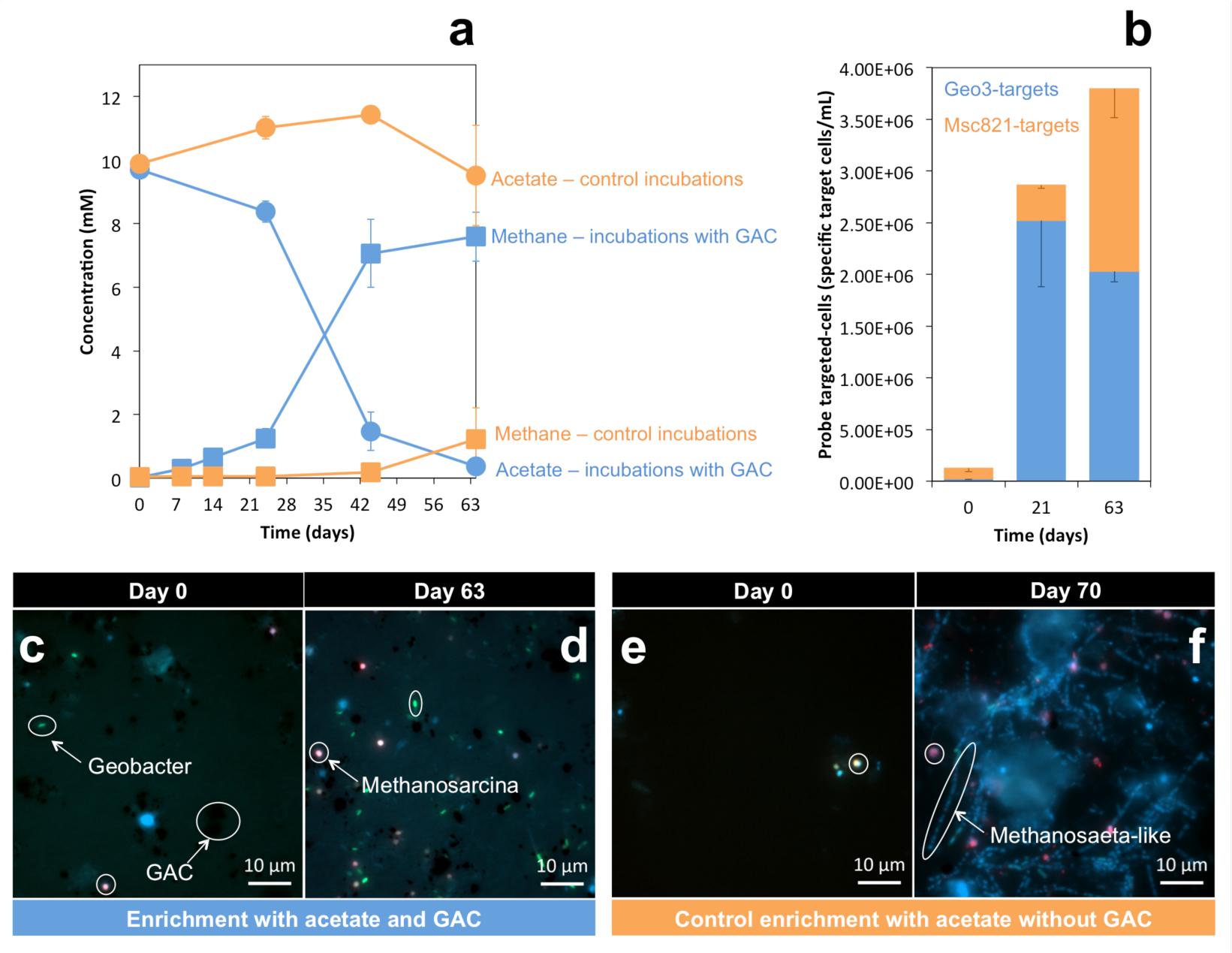
Incubations with and without activated carbon. Acetate consumption and methane production in mud-free slurries provided with (blue lines) and without (orange lines) GAC (a). CARD-FISH with specific probes showed an increase in *Geobacter* cells (Alexa488 tyramides and HRP-labeled Geo3-probe) over time, followed by an increase in *Methanosarcina* cells (Alexa594 and HRP-labeled Msc821 probe) in cultures with conductive activated carbon (c, d). On the other hand, in control cultures without GAC, *Methanosarcina* and *Methanothrix*-like cells were detected, but no *Geobacter* (e, f). Panels c and e show distribution of cells immediately after inoculation, whereas panels d and f show the distribution after 63 and 70 days of incubation, in stationary or late exponential phase, respectively.

### Syntrophic acetate oxidation mediated by GAC

Repeated transfers of the SAO cultures with acetate as electron donor, CO_2_ as electron acceptor and GAC produced methane much faster than GAC-free controls and led to sediment-free cultures enriched in *Geobacter* and *Methanosarcina* (Fig. 3). The enriched *Geobacters* were apparently a new species, 96% related to *G. psychrophilus* (Fig. 2b), whereas *Methanosarcina* were 99% related to *M. subterranea* (Fig. 2c). In the absence of conductive minerals *Geobacter* became depleted already in slurry incubations, while *Methanothrix*-like cells took over acetate-only incubations (Fig. 3f, Supl.file2).

In incubations with acetate and GAC, acetate could be consumed both by acetoclastic methanogens and/or SAO consortia. A schematic representation of SAO mediated by GAC tied to methanogenesis is presented in Fig. 4a. Our hypothesis was that during SAO *Geobacter* cells donate electrons from the oxidation of acetate to GAC, which then plays the role of a transient electron acceptor. Then *Methanosarcina* cells would retrieve the electrons from GAC in order to reduce CO_2_ to methane.

**Figure 4.**
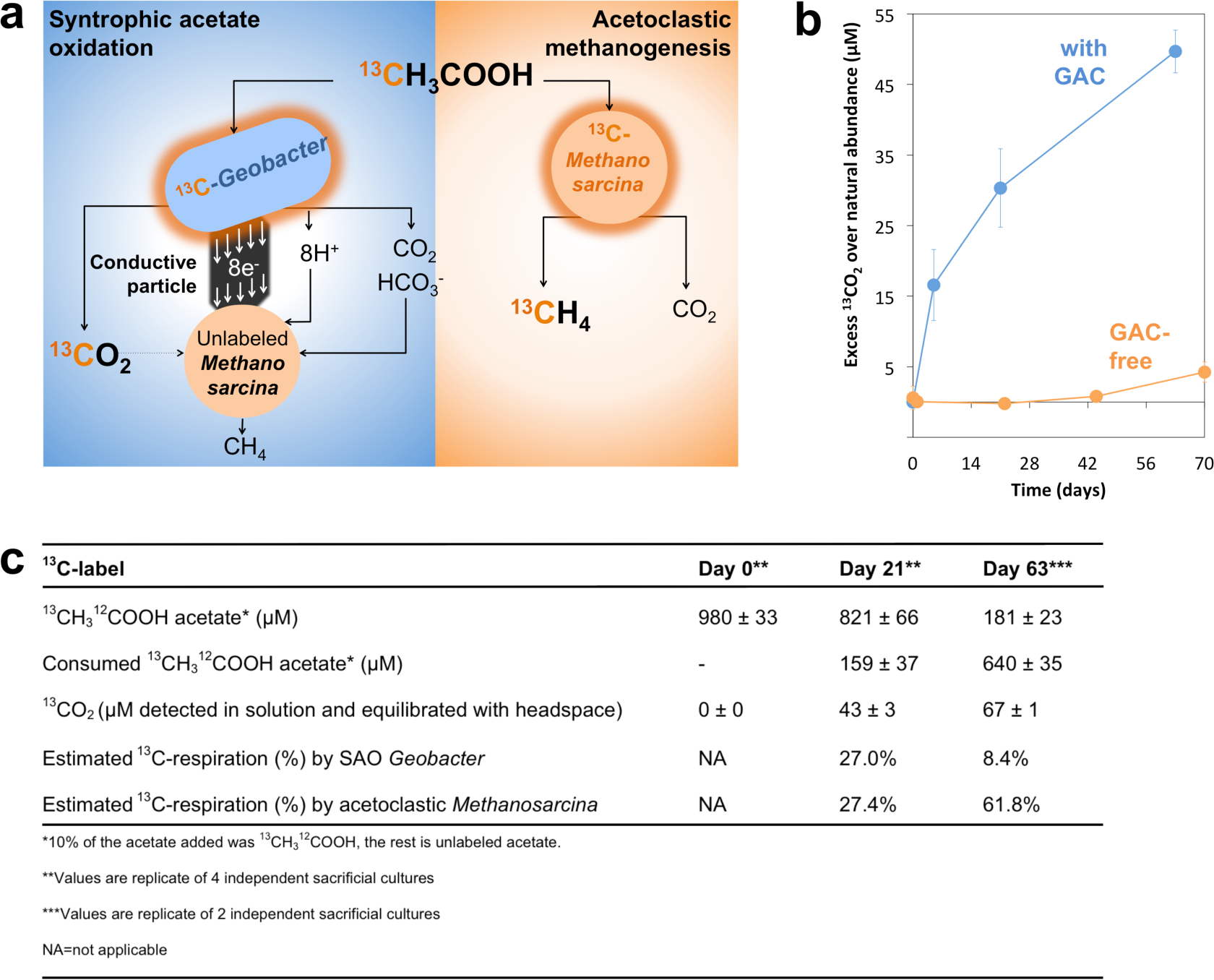
Experimental approach and evidence for SAO. Experimental approach to distinguish between SAO and acetoclastic methanogenesis using isotopic labeling experiments (a). ^13^CH_3_-^12^COOH was provided as 10% of the total acetate, which played the role of the electron donor for SAO-consortia from the Bothnian Bay. During SAO, acetate oxidizing *Geobacter* cells are expected to produce ^13^CO_2_ (^13^C-carbon depicted in orange) and to incorporate ^13^C-acetate. During SAO, ^13^CO_2_ will be diluted by the bicarbonate in the medium and should not generate significant ^13^CH_4_. However, acetoclastic methanogenesis by *Methanosarcina* cells will generate ^13^CH_4_ from ^13^CH_3_^12^COOH while cells incorporate ^13^C-acetate in their cell-mass. Cells expected to incorporate ^13^C-acetate are encircled in orange. SAO activity was validated (b) by labeled ^13^CO_2_ production from acetate especially in SAO-consortia provided with GAC (blue) versus cultures without GAC (orange). An overview of acetate catabolism and how much is used for respiration by *Geobacter* versus acetoclastic methanogenesis by *Methanosarcina* (c).

To distinguish between acetoclastic methanogenesis and SAO, cultures were incubated with ^13^CH_3_^12^COOH. If acetoclastic methanogens utilized the ^13^C-methyl on acetate, they would only produce ^13^CH_4_. However, if SAO-bacteria utilize ^13^C-acetate then they would produce ^13^CO_2_ (Fig.4a). When acetoclastic methanogens and SAO-bacteria use ^13^C-methyl on acetate at the same time, both ^13^CO_2_ and ^13^CH_4_ would be produced. Our results support the later model.

#### SAO dependency on GAC

Incubations for circa 70 days with ^13^C-acetate and GAC converted the ^13^C-methyl on acetate to ^13^CO_2_, whereas control cultures lacking GAC produced little ^13^CO_2_ (Fig. 4). This indicated that indeed GAC stimulated syntrophic acetate oxidation (SAO).

#### Respiratory metabolism and SAO

During exponential growth (day 21) SAO could explain 27% of the total respiratory metabolism whereas 27.4% could be explained by acetoclastic methanogenesis (Fig. 4c). During stationary phase (day 63), SAO justified 8.4% of the total respiratory metabolism whereas acetoclastic methanogenesis justified 61.8%.

#### Biosynthetic metabolism and SAO

The increase in abundance of *Geobacter* cells over time (Fig. 3b) in incubations with GAC indicated that they could play the role of syntrophic acetate oxidizers in SAO consortia. This was confirmed by analysis of the ^13^CH_3_^12^COOH incubated SAO consortia by NanoSIMS/CARD-FISH, an approach that helps correlate phylogeny and function (72). During incubation with GAC both *Geobacter* and *Methanosarcina* cells became greatly enriched in ^13^C, indicating label-assimilation from acetate (Fig. 5a, b). During exponential phase (day 21), *Geobacter* cells were six times more abundant than *Methanosarcina* (Fig. 3b). Therefore, the entire *Geobacter*-population assimilated 5 times more acetate than the *Methanosarcina-*population (Fig. 5a). However, upon prolonged incubation (day 63), the number of *Geobacter* cells remained relatively constant, while *Methanosarcina* cells increased in abundance to match the *Geobacter* population (Fig. 3b). As a consequence, during the late incubation phase, the *Methanosarcina* population assimilated three fold more acetate than *Geobacter* (Fig. 5b).

**Figure 5.**
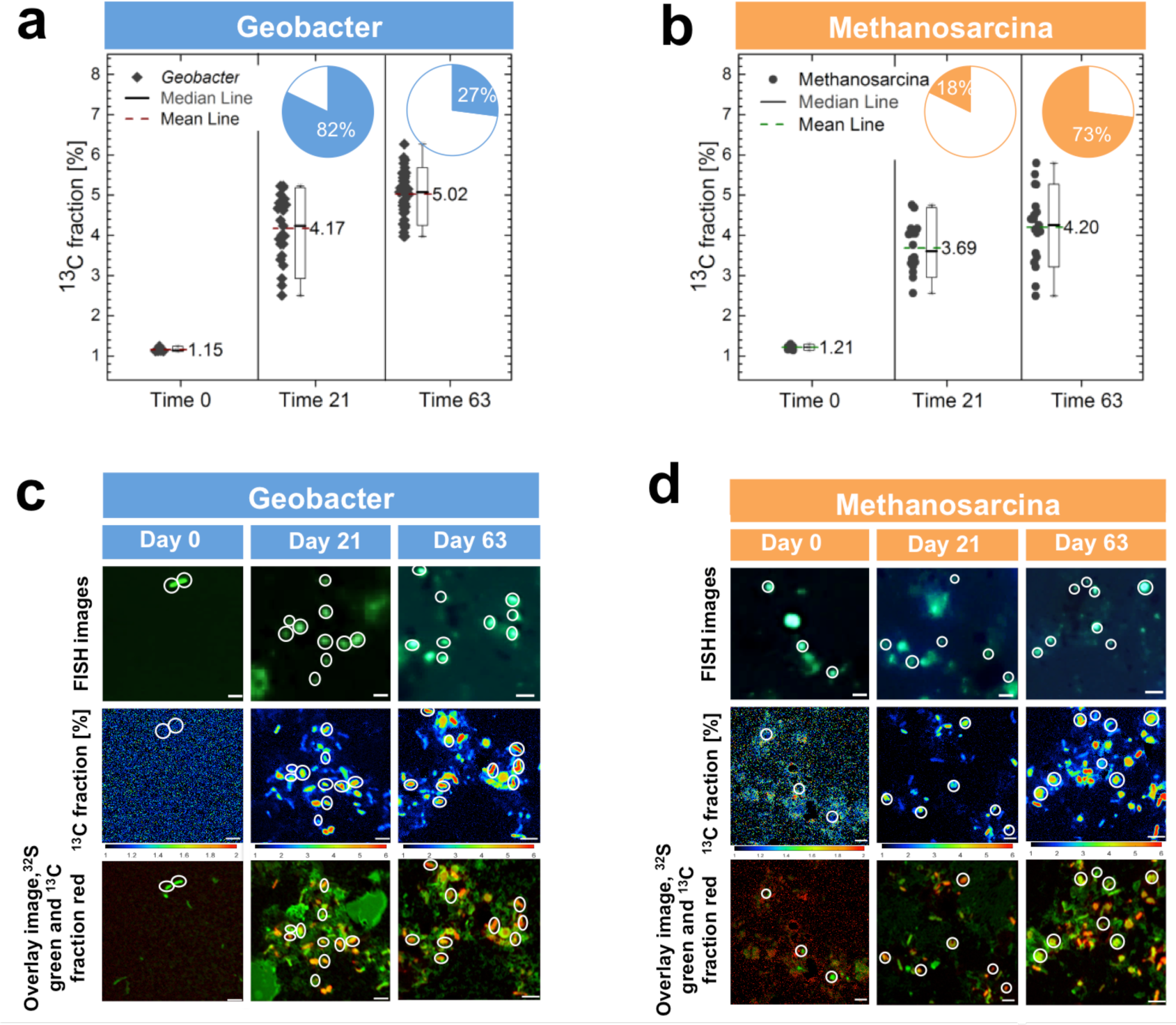
NanoSIMS identification of cells incorporating ^13^C-labeled acetate. Highly abundant *Geobacter* incorporated more ^13^CH_3_-^12^COOH per cell (a) than *Methanosarcina* (b). Insets for panels (a) and (b) show percentage assimilation in *Geobacter* (blue insets) and *Methanosarcina* (orange) over time. (c) Time-dependent distribution of cells labeled by *Geobacter*-specific probes compared with time-dependent incorporation of ^13^CH_3_COOH in *Geobacter*-cells (see scales below images) and an overlay of ^13^C-incorporation (red) to total biomass as detected by tracing ^32^S (green) using NanoSIMS. (d) Time-dependent distribution of cells labeled by *Methanosarcina*-specific probes compared with time-dependent incorporation of ^13^CH_3_COOH in *Methanosarcina*-cells (see scales below images) and an overlay of ^13^C-incorporation (red) to total biomass as detected by tracing ^32^S (green) using NanoSIMS.

The ratio of *Geobacter* to *Methanosarcina* cells in the original sediment (8:1) was more similar to that observed in incubation during exponential growth (6:1) than to that observed during stationary phase (1:1). During exponential growth *Geobacter* cells incorporate high ^13^C-label and their distribution is similar to that from environmental samples. This suggests that *Geobacter* is the primary acetate oxidizer in SAO consortia from the Baltic Sea (Fig. 5).

#### SAO is not decoupled from methanogenesis

The faster growth of *Geobacter* and slower growth of *Methanosarcina* (Fig. 3) implies that *Geobacter* might oxidize acetate alone using GAC as terminal electron acceptor rather than the methanogen. To verify this hypothesis, we blocked the metabolic activity of the methanogen by using a methyl-coenzyme M analogue (10 µM 2-bromoethanesulfonate or BES) to chemically inhibit methanogenesis (67). If *Geobacter* respired GAC, independent of electron uptake by *Methanosarcina*, we should be able to decouple acetate utilization by *Geobacter* from methanogenesis. However, acetate utilization ceased as soon as methanogenesis was inhibited by BES (Fig 6), indicating a dependence of the (exo)electrogenic syntrophic acetate oxidizer (*Geobacter*) on the *Methanosarcinales*-methanogen. We therefore conclude that an acetate oxidizing *Geobacter*, donates electrons to GAC, from where *Methanosarcina* retrieves electrons to reduce CO_2_ to methane.

**Figure 6.**
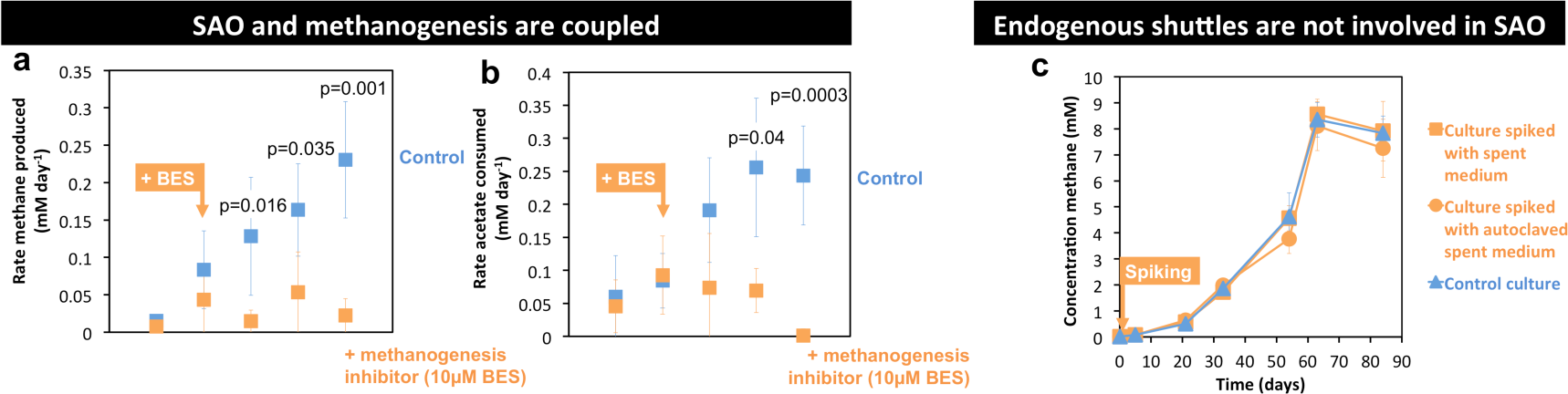
Bothnian Bay enrichments coupled SAO and methanogenesis without the need for inherent soluble shuttles. The methanogenic inhibitor bromoethane sulfonate (BES) collapsed the rates of both methanogenesis (a) and acetate oxidation (b), indicating that the two processes are coupled, and that *Geobacter* cannot grow alone on acetate and GAC. Methane production (a) and acetate utilization (b) rates were measured in cultures spiked with BES (orange), in contrast to controls lacking BES (blue). Mineral-SAO incubations spiked with raw spent media, which was filtered or autoclaved, showed similar methane production as control mineral-SAO incubations without spent media.

#### Exoenzymes and shuttles are not endogenously created

Previous studies indicated that extracellular viable enzymes could act as manufacturers of diffusible chemicals (H_2_, formate) which could be used for electron transfer to methanogens (68). To verify this hypothesis we spiked cultures with spent media from a fully-grown culture, filtered through a 0.2 µm filter. The spent medium should theoretically contain (exo)cellular enzymes or potential shuttles, and if these were involved in electron transfer between the microorganisms from the Bothnian Bay sediments we should see an increase in methanogenic rates. We did not notice an increase in methanogenic rates in spiked cultures compared to control cultures (Fig. 6). This indicates that (exo)cellular enzymes/shuttles were unlikely to play a role in conductive particle mediated SAO between *Geobacter* and *Methanosarcina*.

## Conclusion

Here we showed that syntrophic acetate oxidation was coupled to CO_2_-reductive methanogenesis via conductive materials. Our results demonstrate that conductive particles could be fundamental for syntrophic acetate oxidation coupled to CO_2_-reductive methanogenesis in sediments. We propose that mineral-SAO could have significant implications for the isotopic composition and the cycling of methane in aquatic sediments. Anthropogenic activity could enhance the input of conductive materials to sediments, ultimately increasing methane fluxes. Since methane is a powerful greenhouse gas we must better understand such actuators of methane emissions in the environment.

## Materials and methods

### Sampling and incubations

During an expedition on board of RV Fyrbygarrren in July 2014, we sampled sediment cores with a Gemini gravity corer. Three sediment cores were gathered at station RA2, which is located near the Swedish shoreline (coordinates: 22° 26.8′ East; 65° 43.8′ North). Within 24 hours after sampling, the sediment was partitioned in depth-profiled aliquots and fixed for biogeochemical, and molecular analyses inside an on deck N_2_-inflatable glove bag, as described below in detail.

For incubations, we gathered methanogenic sediment from 30–36 cm depth, and replaced the gas atmosphere with 2 bars N_2_:CO_2_ 80:20 mix. The 30–36 cm depth sediment was stored at 4°C until we generated slurries with various substrates and minerals.

Slurries were prepared in the lab in an anaerobic chamber, and were generated within six months after sampling. For slurries, we used 3 mL cut-off syringes to distribute 2.5 mL sediment mix into 20 mL gas-tight vials filed with 7.5 mL DSM 120 modified media. The modified DSM 120 media was prepared as before (48) but with 0.6 g NaCl. Sediment-slurries had a high organic content, whereas mud-free enrichments did not. Therefore we amended the mud-free enrichments with 0.2 g/L yeast extract from a 100 g/L anaerobic and sterile stock, which is required for methanogenic growth. Before inoculation, the complete media which lacked the substrate and (semi)conductive minerals was dispensed anaerobically by syringe into sterile degased vials with or without minerals prepared as below.

Conductive materials, GAC (0.1 g/10mL; Merck) or magnetite (0.1 g/10mL; Sigma-Aldrich) were weighted, added to vials, overlaid with 200 µl ultrapure water for wet sterilization, degased for 3 minutes with N_2_:CO_2_ 80:20 mix, and autoclaved at 121°C for 25 min. Control experiments were carried out with acid washed glass beads instead of conductive minerals. Substrates (5 mM glucose, 5 mM butyrate, 10 mM acetate, 10 mM ethanol) were added to media from sterile anoxic 1M stocks using aseptic and anaerobic techniques. Control experiments were carried out without additional substrate to learn if the organic in sediment could be used as substrates for methanogenesis. All incubations were carried out at room temperature (20–23°C) in triplicate unless otherwise noted.

Gas samples were withdrawn at timed intervals using hypodermic needled connected to a syringe closed by an airtight valve. Gas samples (0.5 mL) were stored until measured, by displacing 0.5 mL ultrapure water, which filled 3 mL exetainers. 30 µl gas sample was tested for methane on a Thermo Scientific gas chromatograph equipped with a TG-Bond Msieve 5A column (30 m × 0.53 mm × 50 µm), and an FID detector. The carrier was N_2_ (flow 5 mL/min) and we used an isothermal oven temperature of 150°C with the injector and detector set at 200°C. Gas standards (0.01% to 50% CH_4_ in N_2_) from Mikrolab Aarhus A/S were always run along with samples. Short chain volatile fatty acids (SCVFA) were detected by high performance liquid chromatography (HPLC) of 0.45 µm filtered and 3 times diluted samples. For HPLC we used an Agilent 1100 instrument equipped with an Aminex-HPX 87H column heated at 70°C, and a VWR detector which detects SCVFA at 210 nm. 5 mM sulfuric acid was used as eluent at a flow of 0.6 mL/min. Standards used ranged between 0.1 mM and 10 mM. Detection limit for all SCVFA was 100 µM.

### Biogeochemical analyses

To determine biogeochemical parameters, we took sediment aliquots every 2 cm in an anaerobic glove bag filled with N_2_ gas. At this station the sulfide-methane transition zone was below 15 cm. Geochemical parameters of direct relevance to this work were methane, dissolved inorganic carbon (DIC), and resident iron and manganese oxide species. For *in situ* methane concentrations and ^13^C/^12^C-methane isotopic fractionation, we blocked the activity of the microorganisms by immersing 2 mL active sediment into 4 mL NaOH 2.5%. NaOH-treated samples kept in gas-tight vials were stored at 4°C, upside down until methane could be measured.

Methane headspace concentrations were measured on a Perkin Elmer GC equipped with an EliteQPlus capillary column with inner diameter of 0.52 mm heated to 50°C, and an FID detector heated to 200°C. The carrier gas was N_2_ with a flow of 10 mL/min. δ^13^C_CH4_ values were measured at Aarhus University on an isotope-mass ratio IR-GCMS as described before (83).

For determination of insoluble iron and manganese, 5 mL sediment was subsampled from each 2 cm-depth interval, transferred into 15 mL centrifugation vials and stored at −20 °C until extraction of different iron and manganese phases. Three different extraction methods were applied: the cold 0.5 N HCl extraction (to dissolve poorly crystalline iron oxides, FeS and FeCO_3_) the dithionite extraction (to dissolve all the other Fe-oxides except for magnetite) and oxalate extraction (to dissolve magnetite) (69, 70), followed by a ferrozine assay (71). For analysis of insoluble manganese, extractions were carried out as described for solid iron and concentrations in the supernatant were analyzed undiluted by flame atomic absorption spectroscopy.

For porewater parameters, porosity of the sediments was calculated from identifying the relationship between the wet weight of the sediment and its dry weight. For porewater extraction 50 mL sediment was sampled every 2 cm, by scooping sediment in Falcon tubes, from which porewater was extracted with the use of rhizons (Rhizosphere; pore size = 0.2 µm). Porewater work was carried under a N_2_ atmosphere in a glove bag.

For porewater Fe^2+^ and Mn^2+^ concentrations, 1 mL porewater was mixed with 20 µL 6 N HCL and stored at −20°C. Soluble Fe^2+^ in the porewater was determined using the ferrozine assay (71).

Porewater DIC was sampled inside an N_2_-filled glove bag on board. DIC samples were filled to brim to ensure no gas bubbles into 3 mL glass vials, which contained 20 µL HgCl_2_-saturated water. Samples were stored upside down at 4°C until measurements. For measurements, we converted DIC to CO_2_ by acidification with 50 µL undiluted H_2_PO_4_ for each 200 µL DIC sample. CO_2_ was allowed to equilibrate in headspace over night inside 12 mL He-flushed exetainers. DIC concentration and the ^13^C/^12^C-DIC isotope ratios were measured on an isotope ratio mass spectrometer coupled to a gas bench, as previously described (72).

## Molecular analyses

For molecular analyses we sampled 2 mL every 2 cm of sediment depth. Samples were collected using cut-off syringes at the same time with samples for biogeochemical parameters, so on board and inside an anaerobic bag. For safe storage during transportation, 3 depths, so a total of 6 cm, were pooled together and mixed with 6 mL MoBio RNA*later* (MoBio; 1:1). Prior to DNA extractions the RNA*later* was removed by centrifugation. For DNA extraction we used the MoBio RNA Soil kit coupled to a complementary DNA Soil kit and followed the instructions provided by the kit manufacturer. DNA was quantified using a Nano Drop, before downstream applications.

### Quantitative PCR

To target electrogenic microorganisms, genus/order-specific PCR was performed with primers for *Desulfuromonadales* (includes all *Geobacter*), *Geothrix*, *Rhodoferrax* and *Shewanella*. For methanogens the following genus/order specific primers were tested to target: *Methanosarcinaceae*, *Methanothrix*, *Methanococcales, Methanobacteriales*, *Methanomicrobiales*. A list of all the primers used, making of standards and the conditions for qPCR are available in Suppl.file1, respectively.

### 16S rRNA gene sequencing, library preparation and phylogenetic tree reconstruction

16S rRNA gene MiSeq amplicon sequencing was carried out from the 30–36 cm depth interval of triplicate cores. Details on procedure can be found in Suppl.file1. Amplification of partial *Geobacter* and *Methanosarcina* 16S rDNA sequences was done as before (76). Cloning employed the TOPO TA Cloning Kit (ThermoFisher Scientific) followed by direct sequencing of PCR products from cloned plasmid DNA (Macrogen). Maximum likelihood phylogenetic trees were constructed using Geneious (77). Sequence files can be found at NCBI under Bioproject ID: PRJNA415800.

### ^13^C labeling experiments

Cultures were incubated with a mix of 1 to 9 of ^13^CH_3_COOH to unlabeled acetate. Circa 21 cultures with GAC and 16 for the GAC-free cultures were started for the NanoSIMS experiment because we would sacrificially harvest three at each time point. Headspace gas samples and VFA samples were analyzed as above.

We followed enrichment of ^13^CO_2_ over time by IRMS. Briefly, 2.5 mL media samples were retrieved anaerobically for ^13^CO_2_ analyses and immediately stored with 20 µl HgCl_2_-saturated water, without any headspace, acidified like explained above for DIC analyses in sediment samples, and finally IRMS analyses were carried out manually against CO_2_ gas standards and bicarbonate standards.

We followed the incorporation of labeled acetate (^13^CH_3_COOH) into a specific phylotype using CARD-FISH coupled to NanoSIMS, as described below (78).

### CARD FISH

To count cells of a specific phylogenetic group, and label cells prior to NanoSIMS we used catalyzed reporter deposition – fluorescence in situ hybridization (CARD-FISH) as described previously (79). CARD-FISH analysis was performed like before (79) using the following probes: Non338 (80) to check for non-specific binding, Eub338I-III (81, 82) to target *Eubacteria*, Geo3a-c in equimolar amounts with helpers H-Geo3-3, H-Geo3-4 to target the *Geobacterales* cluster (83); Arch915 (84) to target *Archaea*, and MS821 (84) to target *Methanosarcina* species. A detailed description of the CARD FISH protocol can be found in Suppl.file2.

### Quantitative imaging of ^13^C label incorporation by NANO SIMS

Chemical imaging and quantitative analysis of ^13^C label incorporation was carried out on a nano-focused Secondary Ion Mass Spectrometry (NanoSIMS-50L) instrument (CAMECA, AMETEK) operating in negative extraction mode. NanoSIMS analyses were carried out on laser micro-dissection selected fields and the collected data were quantitatively analyzed using the LANS software (85). A detailed description of the protocol used for NanoSIMS analyses and data collection can be found in Suppl.file2.

## Acknowledgements

This work is a contribution to Danish Research Council (DFF) grant 1325-00022 to AER. DFF grant 4181-00203 supported OSW. The RV Fyrbyggaren expedition was co-financed a Swedish Research Council (VR) grant to PH and a DFF grant 4002-00521 to BT. We acknowledge the Centre for Chemical Microscopy (ProVIS) at the Helmholtz Centre for Environmental Research supported by the European Regional Development Funds (EFRE – Europe funds Saxony) for using their analytical facilities. We would like to thank Lasse Ørum Smidt, Heidi Grøn Jensen, Susanne Møller, Erik Laursen and Karina Henricksen, Dr. Laura Bristow, and Prof. Dr. Fanghua Liu for help with different aspects of the work.

## Contribution

Experimental design by AER, BOT and NM; Sampling by AER, POH and HSW; Incubations by AER; Biogeochemical parameters by AER, POH, BOT and HSW; Community analyses by AER, PMS, OSW; Stable isotope experiments and analyses by AER, BOT and FM; CARD-FISH coupled with NanoSIMS experiments and analyses by AER, FC, HS, HHR, FM, and NM; Manuscript written by AER with contribution from all authors.

## References

1. Hattori S (2008) Minireview Syntrophic Acetate-Oxidizing Microbes in Methanogenic Environments. Microbes Environ 23(2):118–127.

2. Galushko AS, Schink B (2000) Oxidation of acetate through reactions of the citric acid cycle by Geobacter sulfurreducens in pure culture and in syntrophic coculture. Arch Microbiol 174(5):314–321.

3. Hattori S, Kamagata Y, Hanada S (2000) Thermacetogenium phaeum gen. nov., sp. nov., a strictly anaerobic, thermophilic, syntrophic acetate-oxidizing bacterium. Int J Syst Evol Microbiol 50(2000):1601–1609.

4. Schnurer A, Schink B, Svensson BOH (1996) Clostridium ultunense sp. nov., a mesophilic bacterium oxidizing acetate in syntrophic association with a hydrogenotrophic methanogenic bacterium. Int J Syst Bacteriol 46(4).

5. Balk M, Weijma J, Stams AJM (2002) Thermotoga lettingae sp. nov., a novel thermophilic, methanol-degrading bacterium isolated from a thermophilic anaerobic reactor. Int J Syst Evol Microbiol 52(4):1361–1368.

6. Westerholm M, Roos S, Schnürer A (2010) Syntrophaceticus schinkii gen. nov., sp. nov., an anaerobic, syntrophic acetate-oxidizing bacterium isolated from a mesophilic anaerobic filter. FEMS Microbiol Lett 309(1):100–104.

7. Zhilina TN, Zavarzina DG, Kolganova T V., Tourova TP, Zavarzin GA (2005) "Candidatus Contubernalis alkalaceticum,” an obligately syntrophic alkaliphilic bacterium capable of anaerobic acetate oxidation in a coculture with Desulfonatronum cooperativum. Microbiology 74(6):695–703.

8. Cord-Ruwisch R, Lovley DR, Schink B (1998) Growth of Geobacter sulfurreducens with acetate in syntrophic cooperation with hydrogen-oxidizing anaerobic partners. App/ Environ Microbio/ 64(6):2232–2236.

9. Kimura Z, Okabe S (2013) Acetate oxidation by syntrophic association between Geobacter sulfurreducens and a hydrogen-utilizing exoelectrogen. ISME J 7(8):1472–82.

10. Wang L, Nevin KP, Woodard TL, Mu B, Lovley DR (2016) Expanding the diet for DIET: electron donors supporting direct interspecies electron transfer (DIET) in defined co-cultures. Front Microbiol 7(March):1–7.

11. Galouchko AS, Rozanova EP (1996) Sulfidogenic oxidation of acetate by a syntrophic association of anaerobic mesophilic bacteria. Microbiology 65(February):134–139.

12. Tang J, Zhuang L, Tang Z, Yu Z, Zhou S (2016) Secondary mineralization of ferrihydrite affects microbial methanogenesis in Geobacter - Methanosarcina cocultures. App/ Environ Microbio/ 82(19):5869–5877.

13. Zinder SH (1989) Syntrophic acetate oxidation and “reversible acetogenesis.” Acetogenesis, ed Drake HL (Chapman and Hall).

14. Zinder SH, Koch M (1984) Non-aceticlastic methanogenesis from acetate: acetate oxidation by a thermophilic syntrophic coculture. Arch Microbio/ 138(3):263–272.

15. Phelps TJ, Conrad R, Zeikus JG (1985) Sulfate-dependent interspecies H2 transfer between Methnosarcina barkeri and Desulfovibrio vulgaris during coculture metabolism of acetate or methanol. App/ Environ Microbio/ 50(3):589–594.

16. Ozuolmez D, et al. (2015) Methanogenic archaea and sulfate reducing bacteria co-cultured on acetate: Teamwork or coexistence? Front Microbio/ 6(MAY):1–12.

17. Petersen SP, Ahring BK (1991) Acetate oxidation in a thermophilic anaerobic sewage-sludge digestor: the importance of non-aceticlastic methanogenesis from acetate. FEMS Microbio/ Lett 86(2):149–158.

18. Schnurer A, Zellner G, Svensson BH (1999) Mesophilic syntrophic acetate oxidation during methane formation in biogas reactors. FEMS Microbiol Ecol 29:249–261.

19. Lee SH, et al. (2015) Evidence of syntrophic acetate oxidation by Spirochaetes during anaerobic methane production. Bioresour Technol 190:543–549.

20. Angenent LT, Sung S, Raskin L (2002) Methanogenic population dynamics during startup of a full-scale anaerobic sequencing batch reactor treating swine waste. Water Res 36(18):4648–4654.

21. Karakashev D, Batstone DJ, Trably E, Angelidaki I (2006) Acetate oxidation is the dominant methanogenic pathway from acetate in the absence of Methanosaetaceae. Appl Environ Microbiol 72(7):5138–5141.

22. Tatara M, Makiuchi T, Ueno Y, Goto M, Sode K (2008) Methanogenesis from acetate and propionate by thermophilic down-flow anaerobic packed-bed reactor. Bioresour Technol 99(11):4786–4795.

23. Westerholm M, et al. (2011) Quantification of syntrophic acetate-oxidizing microbial communities in biogas processes. Environ Microbiol Rep 3(4):500–505.

24. Ho D, et al. (2016) High-rate, high temperature acetotrophic methanogenesis governed by a three population consortium in anaerobic bioreactors. PLoS One 11(8): 1–13.

25. Mosbæk F, et al. (2016) Identification of syntrophic acetate-oxidizing bacteria in anaerobic digesters. ISME J 2:1–14.

26. Müller B, Sun L, Westerholm M, Schnürer A (2016) Bacterial community composition and fhs profiles of low - and high - ammonia biogas digesters reveal novel syntrophic acetate - oxidising bacteria. Biotechnol Biofuels: 1–18.

27. Nüsslein B, Chin KJ, Eckert W, Conrad R (2001) Evidence for anaerobic syntrophic acetate oxidation during methane production in the profundal sediment of subtropical Lake Kinneret (Israel). Environ Microbiol 3(7):460–470.

28. Zheng S, et al. (2015) Co-occurrence of Methanosarcina mazei and Geobacteraceae in an iron(III)-reducing enrichment culture. Front Microbiol 6. doi:10.3389/fmicb.2015.00941.

29. Chauhan A, Ogram A (2006) Fatty acid-oxidizing consortia along a nutrient gradient in the Florida Everglades. Appl Environ Microbiol 72(4):2400–2406.

30. Hori T, Noll M, Igarashi Y, Friedrich MW, Conrad R (2007) Identification of acetate-assimilating microorganisms under methanogenic conditions in anoxic rice field soil by comparative stable isotope probing of RNA. Appl Environ Microbiol 73(1):101–109.

31. Liu F, Conrad R (2010) Thermoanaerobacteriaceae oxidize acetate in methanogenic rice field soil at 50°C. Environ Microbiol 12(8):2341–2354.

32. Rui J, Qiu Q, Lu Y (2011) Syntrophic acetate oxidation under thermophilic methanogenic conditions in Chinese paddy field soil. FEMS Microbiol Ecol 77:264–273.

33. Mayumi D, et al. (2011) Evidence for syntrophic acetate oxidation coupled to hydrogenotrophic methanogenesis in the high-temperature petroleum reservoir of Yabase oil field (Japan). Environ Microbiol 13(8):1995–2006.

34. Jensen MM, Thamdrup B, Rysgaard S, Holmer M, Fossing H (2003) Rates and regulation of microbial iron reduction in sediments of the Baltic-North Sea transition. Biogeochemistry 65(3):295–317.

35. Maher B, Taylor R (1988) Formation of ultrafine-grained magnetite in soils. Nature 336(6197):368–370.

36. Nielsen LP, Risgaard-Petersen N, Fossing H, Christensen PB, Sayama M (2010) Electric currents couple spatially separated biogeochemical processes in marine sediment. Nature 463(7284):1071–1074.

37. Liesack W, Schnell S, Revsbech NP (2000) Microbiology of flooded rice paddies. FEMS Microbiol Rev 24(5). doi:10.1016/S0168-6445(00)00050-4.

38. Schmidt MWI, Noack AG (2000) Black carbon in soils and sediments:Analysis, distribution, implications, and current challenges. Global Biogeochem Cycles 14(3):777–793.

39. Middelburg JJ, Nieuwenhuize J, Breugel P van (1999) Black carbon in marine sediments. Mar Chem 65(September 1998):245–252.

40. Marinho B, Ghislandi M, Tkalya E, Koning CE, de With G (2012) Electrical conductivity of compacts of graphene, multi-wall carbon nanotubes, carbon black, and graphite powder. Powder Technol 221:351–358.

41. Liu F, et al. (2015) Magnetite compensates for the lack of a pilin-associated c-type cytochrome in extracellular electron exchange. Environ Microbiol 17(3):648–55.

42. Liu F, et al. (2012) Promoting direct interspecies electron transfer with activated carbon. Energy Environ Sci 5(10):8982.

43. Chen S, et al. (2014) Promoting interspecies electron transfer with biochar. Sci Rep 4:5019.

44. Chen S, et al. (2014) Carbon cloth stimulates direct interspecies electron transfer in syntrophic co-cultures. Bioresour Technol 173:82–6.

45. Yang Z, Shi X, Wang C, Wang L, Guo R (2015) Magnetite nanoparticles facilitate methane production from ethanol via acting as electron acceptors. Sci Rep 5:16118.

46. Zhuang L, Tang J, Wang Y, Hu M, Zhou S (2015) Conductive iron oxide minerals accelerate syntrophic cooperation in methanogenic benzoate degradation. J Hazard Mater 293(808):37–45.

47. Summers ZM, et al. (2010) Direct exchange of electrons within aggregates of an evolved syntrophic coculture of anaerobic bacteria. Science (80-) 330(6009):1413–1415.

48. Rotaru A-E, et al. (2014) Direct interspecies electron transfer between Geobacter metallireducens and Methanosarcina barkeri. Appl Environ Microbiol 80(15):4599–605.

49. Rotaru A-E, et al. (2014) A new model for electron flow during anaerobic digestion: direct interspecies electron transfer to Methanosaeta for the reduction of carbon dioxide to methane. Energy Environ Sci 7(1):408.

50. Zhao Z, Zhang Y, Woodard TL, Nevin KP, Lovley DR (2015) Enhancing syntrophic metabolism in up-flow anaerobic sludge blanket reactors with conductive carbon materials. Bioresour Techno/ 191:140–145.

51. Xu S, et al. (2015) Comparing activated carbon of different particle sizes on enhancing methane generation in upflow anaerobic digester. Bioresour Techno/ 196:606–612.

52. Cruz Viggi C, et al. (2014) Magnetite particles triggering a faster and more robust syntrophic pathway of methanogenic propionate degradation. Environ Sci Techno/ 48(13):7536–7543.

53. Li H, et al. (2015) Direct interspecies electron transfer accelerates syntrophic oxidation of butyrate in paddy soil enrichments. Environ Microbio/ 17(5):1533–1547.

54. Zhang J, Lu Y (2016) Conductive Fe3O4 nanoparticles accelerate syntrophic methane production from butyrate oxidation in two different lake sediments. Front Microbio/ 7(AUG):1–9.

55. Dolfing J (2014) Thermodynamic constraints on syntrophic acetate oxidation. App/ Environ Microbio/ 80(4):1539–1541.

56. Kato S, Hashimoto K, Watanabe K (2012) Microbial interspecies electron transfer via electric currents through conductive minerals. Proc Nat/ Acad Sci 109(25):10042–10046.

57. Conrad R, Klose M (2011) Stable carbon isotope discrimination in rice field soil during acetate turnover by syntrophic acetate oxidation or acetoclastic methanogenesis. Geochim Cosmochim Acta 75(6):1531–1539.

58. Kastening B, Hahn M, Rabanus B, Heins M, Felde U (1997) Electronic properties and double layer of activated carbon? 42(18).

59. Fallas Dotti M (2015) Origin and evolition of sulfur processing organisms through time. Dissertation (University of Southern Denmark).

60. Egger M, et al. (2014) Iron-mediated anaerobic oxidation of methane in brackish coastal sediments. Environ Sci Technol 277(1). doi:10.1021/es503663z.

61. Sanchez-Garcia L, Cato I, Gustafsson Ö (2010) Evaluation of the influence of black carbon on the distribution of PAHs in sediments from along the entire Swedish continental shelf. Mar Chem 119(1–4):44–51.

62. Edlund A, Hårdeman F, Jansson JK, Sjöling S (2008) Active bacterial community structure along vertical redox gradients in Baltic Sea sediment. Environ Microbiol 10(8):2051–2063.

63. Fey A, Claus P, Conrad R (2004) Temporal change of 13C-isotope signatures and methanogenic pathways in rice field soil incubated anoxically at different temperatures. Geochim Cosmochim Acta 68(2):293–306.

64. Rotaru A-E, Woodard TL, Nevin KP, Lovley DR (2015) Link between capacity for current production and syntrophic growth in Geobacter species. Front Microbiol 6:744.

65. Thauer RK, Kaster A-K, Seedorf H, Buckel W, Hedderich R (2008) Methanogenic archaea: ecologically relevant differences in energy conservation. Nat Rev Microbiol 6(8):579–91.

66. Roden EE, Zachara JM (1996) Microbial reduction of crystalline iron (III) oxides: influence of oxide surface area and potential for cell growth. Environ Sci Technol Sci Technol 30(5):1618–1628.

67. Van Bodegom PM, Scholten JCM, Stams AJM (2004) Direct inhibition of methanogenesis by ferric iron. FEMS Microbiol Ecol 49(2):261–268.

68. Deutzmann JS, Sahin M, Spormann AM (2015) Extracellular enzymes facilitate electron uptake in biocorrosion and bioelectrosynthesis. MBio 6(2):e00496–15.

69. Thamdrup B, Fossing H, Jørgensen BB (1994) Manganese, iron and sulfur cycling in a coastal marine sediment, Aarhus Bay, Denmark. Geochim Cosmochim Acta 58:5115–5129.

70. Kostka JE, Luther GW (1994) Partitioning and speciation of solid phase iron in saltmarsh sediments. Geochim Cosmochim Acta 58(7):1701–1710.

71. Lovley DR, Phillips EJP (1987) Rapid Assay for Microbially Reducible Ferric Iron in Aquatic Sediments. 53(7):1536–1540.

72. Nordi KÁ, Thamdrup B, Schubert CJ (2013) Anaerobic oxidation of methane in an iron-rich Danish freshwater lake sediment. Limnol Oceanogr 58(2):546–554.

73. Lloyd KG, MacGregor BJ, Teske A (2010) Quantitative PCR methods for RNA and DNA in marine sediments: Maximizing yield while overcoming inhibition. FEMS Microbiol Ecol 72(1):143–151.

74. Klindworth A, et al. (2013) Evaluation of general 16S ribosomal RNA gene PCR primers for classical and next-generation sequencing-based diversity studies. Nucleic Acids Res 41(1): 1–11.

75. Li W, Fu L, Niu B, Wu S, Wooley J (2012) Ultrafast clustering algorithms for metagenomic sequence analysis. Brief Bioinform 13(6):656–668.

76. Snoeyenbos-West OL, Nevin KP, Anderson RT, Lovley DR (2000) Enrichment of Geobacter species in response to stimulation of Fe(III) reduction in sandy aquifer sediments. Microb Ecol 39(2):153–167.

77. Kearse M, et al. (2012) Geneious Basic: An integrated and extendable desktop software platform for the organization and analysis of sequence data. Bioinformatics 28(12):1647–1649.

78. Musat N, et al. (2008) A single-cell view on the ecophysiology of anaerobic phototrophic bacteria. Proc Natl Acad Sci 105(46):17861–6.

79. Pernthaler J, Glöckner FO, Schönhuber W, Amann R Fluorescence in situ hybridization with rRNA-targeted oligonucleotide probes. Methods in Microv, pp 1–31.

80. Wallner G, Amann R, Beisker W (1993) Optimizing fluorescent in situ hybridization with rRNA-targeted oligonucleotide probes for flow cytometric identification of microorganisms. Cytometry 14(2):136–143.

81. Amann RI, et al. (1990) oligonucleotide probes with flow cytometry for analyzing mixed microbial populations. Combination of 16S rRNA-Targeted Oligonucleotide Probes with Flow Cytometry for Analyzing Mixed Microbial Populations. App/ Environ Microbio/ 56(6):1919–1925.

82. Daims H, Brühl a, Amann R, Schleifer KH, Wagner M (1999) The domain-specific probe EUB338 is insufficient for the detection of all Bacteria: development and evaluation of a more comprehensive probe set. Syst App/ Microbio/ 22(3):434–44.

83. Richter H, Lanthier M, Nevin KP, Lovley DR (2007) Lack of electricity production by Pelobacter carbinolicus indicates that the capacity for Fe(III) oxide reduction does not necessarily confer electron transfer ability to fuel cell anodes. App/ Environ Microbio/ 73(16):5347–5353.

84. Raskin L, Stromley JM, Rittmann BE, Stahl D a (1994) Group-specific 16S rRNA hybridization probes to describe natural communities of methanogens. App/ Environ Microbio/ 60(4):1232–40.

85. Polerecky L, et al. (2012) Look@NanoSIMS - a tool for the analysis of nanoSIMS data in environmental microbiology. Environ Microbio/ 14(4):1009–1023.

86. Games L, Hayes JM (1978) Methane-producing bacteria: natural fractionations of tbe stable carbon isotopes. Geochim Cosmochim Acta 42:1295–1297.

87. Krzycki J, Kenealy W, DeNiro M, Zeikus JG (1987) Stable Carbon Isotope Fractionation by Methanosarcina barkeri during Methanogenesis from Acetate, Methanol, or Carbon Dioxide-Hydrogen. App/ Environ Microbio/ 53(10):2597–2599.

88. Summons RE, Franzmann PD, Nichols PD (1998) Carbon isotopic fractionation associated with methylotrophic methanogenesis. 28(7):465–475.

89. Gelwicks JT, Risatti JB, Hayes JM (1994) Carbon Isotope Effects Associated with Aceticlastic Methanogenesis. 60(2):467–472.

